# The Moral Brain in Context: Political Orientation, Empathy, and Evaluation of Everyday Behavior

**DOI:** 10.1101/2025.08.23.671907

**Authors:** Ronja Demel, Michael R. Waldmann, Anne Schacht

## Abstract

Moral judgments are central to guiding social interactions, influencing whom we trust, cooperate with, or avoid. While extensive research has examined the cognitive and affective bases of moral evaluation, less is known about how stable traits – such as political orientation and empathy – shape these judgments and their neural correlates in ecologically valid contexts. Across two studies, we investigated how people evaluate individuals who committed harmful, helpful, or neutral actions. In Study 1 (N = 214), participants read everyday moral scenarios paired with neutral faces. Empathy moderated gender effects in judging harmful acts, and political orientation selectively influenced ratings of morally ambiguous (neutral) actions. In Study 2 (N = 39), pre-screened conservatives and progressives (N = 675 prescreen) viewed the same agents in an EEG paradigm. Likeability ratings varied by moral valence but not political orientation. Event-related potential analyses revealed that conservatives showed consistently elevated Late Positive Potentials (LPPs) across all conditions, whereas progressives displayed selective LPP enhancement for helpful individuals, suggesting group differences in motivational salience attribution. Together, these findings show that political orientation and empathy influence both explicit moral evaluations and neural processing of socially relevant information. Such insights deepen our understanding of how enduring traits shape moral judgments.

## General Introduction

Accurately evaluating others is fundamental to social interaction. Faces, as primary social stimuli, enable rapid impression formation (Todorov et al., 2007; Willis & Todorov, 2006), but are highly malleable when moral person knowledge is introduced (Barrett et al., 2011; Todorov & Uleman, 2003). Rather than judging individuals solely on the basis of appearance, observers integrate contextual and behavioral cues into person-centered impressions that guide interactions (Albohn et al., 2025).

Morality is central to these evaluations and closely linked to emotion. While early theories emphasized deliberate reasoning in moral judgment (Kohlberg, 1969), dual-process models and affective-neuroscience perspectives highlight the role of intuitive, emotionally grounded processes (Greene et al., 2001; Haidt, 2001). These rapid, emotionally charged judgments help people navigate complex social situations efficiently (Cushman, 2019; Decety & Cowell, 2015).

Empathy – the capacity to share others’ feelings – can heighten moral sensitivity (e.g., Decety & Cowell, 2015; Morelli et al., 2014), but its influence depends on target characteristics, violation type, and context (Bloom, 2016; Koenigs et al., 2007). Higher empathy increases the salience of morally relevant information and may interact with gender, as women often report slightly higher empathy and stronger affective responses to violations (Christov-Moore et al., 2014; Friesdorf et al., 2015).

Political orientation also shapes moral cognition. While progressives and conservatives share core moral concerns, they prioritize different moral foundations (Haidt & Graham, 2007; Graham et al., 2009). Conservatives tend to endorse loyalty, authority, and purity, whereas progressives emphasize care and fairness. Conservatives often show stronger physiological responses to emotionally salient stimuli (Carraro et al., 2011; Oxley et al., 2008; Tritt et al., 2013) and greater affective engagement with taboo violations (Jost et al., 2003; Hibbing et al., 2014). Political orientation can also influence empathy: conservatives have demonstrated more balanced empathy across political groups, while progressives show reduced empathy toward political outgroups (Casey et al., 2023). Furthermore, neural evidence supports ideological differences in empathic processing (Zebarjadi et al., 2023).

Despite such findings, few studies investigated how empathy and political orientation jointly influence moral evaluations, especially using ecologically valid paradigms. Recent work in associative learning has demonstrated that even minimal moral context can profoundly alter face perception (Suess et al., 2014).

Building on this previous research, we conducted two studies testing how political orientation, empathy, and gender shape evaluations of faces in moral contexts. Study 1 (N = 214) examined moral ratings in a large online sample; Study 2 (N = 39) recorded ERPs in a pre-screened sample of conservative and progressive participants using a face–scenario–paradigm. Together, they provide a multilevel perspective on person-centered moral impression formation.

### Study 1 – Moral Impressions from Faces: How Empathy, Gender, and Political Orientation Shape Social Assessments

In daily life, we frequently form rapid impressions of others from minimal cues, such as a face or brief social interaction. Morality is central to these evaluations, guiding decisions about trust, cooperation, and inclusion (Goodwin et al., 2014). Observers often infer enduring traits (e.g., “good” or “bad”) from moral acts (Uhlmann et al., 2015), especially when person-specific features such as faces are available (Todorov & Uleman, 2003; Barrett et al., 2011).

The *dimensional moral model* (Gray & Wegner, 2011) classifies moral events by valence (harm vs. help) and moral type (agent vs. patient). It facilitates systematic study of everyday moral interactions, including morally positive and neutral acts that have often been neglected in moral cognition research (e.g., Clark et al., 2020). This framework offers greater ecological validity than more artificial paradigms like the trolley dilemma (e.g., Clifford et al., 2015; Kahane, 2015).

Faces are potent triggers of moral impressions, particularly when paired with person-specific contextual information (Haxby et al., 2000; Wieser & Brosch, 2012). Even brief verbal associations can imbue neutral faces with valence (Morel et al., 2012; Suess et al., 2014), and such evaluations can be shaped by a single learning episode (Schwarz et al., 2013).

Study 1 tested how individual differences influence person-centered moral evaluations in an ecologically valid paradigm. We presented participants with short, pre-validated scenarios describing helpful, harmful, or neutral acts, each paired with a neutral face and name. This design allowed us to examine how moral context shapes evaluations of specific agents rather than abstract behaviors.

To capture stable traits relevant to moral judgment, participants completed validated measures of empathy and political orientation. Gender was also recorded to assess potential interactions, given prior evidence for gender-linked variability in empathic responses. Moral evaluations were collected on a 7-point scale ranging from “very harmful” to “very helpful.”

This approach offers several advantages. First, the inclusion of morally positive and neutral behaviors provides a broader evaluative range than paradigms focusing solely on moral transgressions. Second, pairing scenarios with identifiable agents ensures person-centered judgments rather than abstract moral reasoning. Third, the simultaneous assessment of empathy, political orientation, and gender allows for the exploration of main and interactive effects within a single model.

#### Hypotheses

1. **Valence** effect: Helpful agents will be rated more positively, and harmful agents more negatively, than neutral agents.
2. **Empathy and gender** will influence sensitivity to moral behavior:

a. Higher empathy will be associated with greater sensitivity to both harmful and helpful behaviors.
b. Female participants will exhibit greater sensitivity to moral behavior compared to male participants
c. Empathy and gender will interact, with empathic women showing the highest sensitivity to harmful and helpful behavior.
3. **Political orientation** will influence moral evaluations: Conservatives will give more extreme ratings (more negative for harm, more positive for help) than progressives.
4. **Trait correlation**: Empathy and political orientation will be positively correlated.

## Methods

### Participants

We recruited 228 German native speakers (18–35 years) via campus and online job boards. Fourteen were excluded because of age or language comprehension issues, leaving 214 participants (115 females, 99 males, *M*_age_ = 23.41, *SD* = 2.75). Sample size was based on prior studies of empathy and political orientation in moral judgment.

### Materials

Stimuli comprised 180 neutral face images from a validated database (Kulke et al., 2017), cropped and luminance-matched. Short moral scenarios (N = 106) were developed based on Gray and Wegner’s (2011) dimensional moral model and pilot-rated (N = 45) for valence, morality, and plausibility. Thirty highly rated scenarios per condition (helpful, harmful, neutral) were selected, each describing intentional acts between an agent and a patient in gender-neutral language (see Table 1 for examples of the scenarios). Only the agent was later evaluated.

**Table 1.** Examples of the scenarios.

|  |  |
| --- | --- |
| Positive | Anna is sitting in a wheelchair that has become stuck in a rut while crossing the train tracks. Jan sees a train approaching in the distance. He runs to Anna on the tracks and brings her to safety. |
| Neutral | Lena and Nico attend the same literature club ever since their first semester. Nico received a book from his grandmother for Christmas. Lena asks him if he liked it. |
| Negative | Hanna asks Anton to take a letter (containing important application documents) to the post office for her. Anton throws the letter in the trash instead of taking it to the post office. |

### Questionnaires

Political orientation was assessed with a German conservatism scale (König & Frank, 2000; *α* = .87). Scores ranged from 2.03–3.71 (*M* = 3.09, *SD* = 0.29), indicating a progressive-leaning sample. Empathy was measured using a German version of the Interpersonal Reactivity Index (SPF; Paulus, 2016; *α* = .80), comprising 12 items on perspective taking, fantasy, and empathic concern.

### Ethics

The study was approved by the ethics committee of the Institute of Psychology, University of Göttingen, and conducted in accordance with the Declaration of Helsinki.

### Procedure

Participants completed the task in view-shielded lab cabins in groups of 4–10. Stimuli were presented using PsychoPy (Peirce, 2007). Each trial began with a fixation cross (1,000 ms), followed by two faces with names (3,000–7,000 ms), and a moral scenario (7,000–20,000s). After another fixation (1,000 ms), the agent’s face (1,000 ms) appeared, and participants rated the agent’s behavior on a 7-point Likert scale (1 = very harmful, 7 = very helpful; see Figure 1). A one-back face memory task occurred randomly in 20% of trials to ensure attention. After the main task, participants completed the questionnaires. Compensation amounted to 5€/hour plus bonus up to 5.40€.

**Figure 1.**
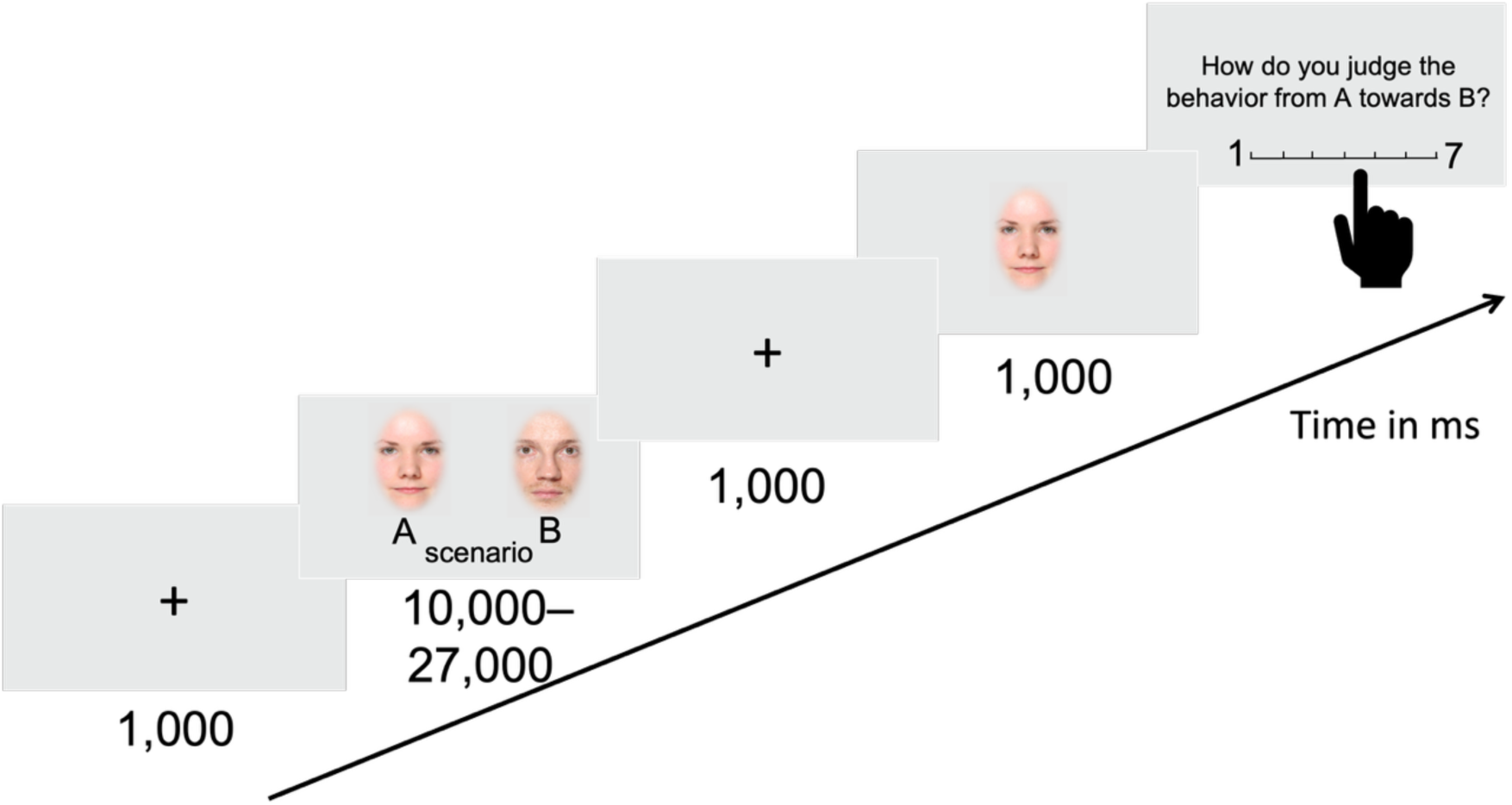
Trial scheme of the task in Study 1.

### Statistical Analyses

Analyses were conducted in R (version 3.5.2; R Core Team, 2019), using linear mixed-effects models (LMMs; lmer function, lme4 package (Bates et al., 2015)). Fixed effects were moral valence (positive, neutral, negative), gender, political orientation, and empathy, with all two-way interactions and a valence × empathy × gender interaction. Participant and item were random intercepts.

Empathy and political orientation scores were z-transformed. Model stability was assessed by iteratively excluding participants and inspecting parameter estimates. Normality and homogeneity were evaluated via residual and QQ plots. Collinearity was checked using the vif function (car package; Fox et al., 2012). Fixed effects were tested by comparing full to null models (likelihood ratio tests; anova, test = “Chisq”), and specific interactions were evaluated with drop1 (Barr, 2013). Significant interactions were explored in valence-specific follow-up models. Pearson correlation (stats package; cor.test) examined empathy-political orientation relationship. The study was not preregistered; full analysis code is available at https://osf.io/qsbvx/?view_only=dd12d36861f240288d3533724fcce665.

## Results

### Descriptive Statistics

Moral ratings (1 = very harmful to 7 = very helpful) varied by scenario valence: harmful scenarios were rated lowest (*M* = 1.61, *SD* = 0.84), neutral scenarios moderate (*M* = 4.62, *SD* = 0.91), and helpful scenarios highest (*M* = 6.39, *SD* = 0.88, see Appendix, Table 1).

### Main Effects and Interactions

The full model was significant compared to the null model, *χ*²(14) = 605.83, *p* < .001, *R²_marginal_* = 0.84, yielding a significant three-way interaction of moral valence × gender × empathy, *χ²*(3) = 63.58, *p* < .001, and a significant moral valence × political orientation interaction, *χ²*(2) = 89.99, *p* < .001 (see Figure 2). Ratings differed significantly between helpful and neutral scenarios (*t* = 17.90, *p* < .001) and between harmful and neutral scenarios (*t* = -30.43, *p* < .001).

**Figure 2.**
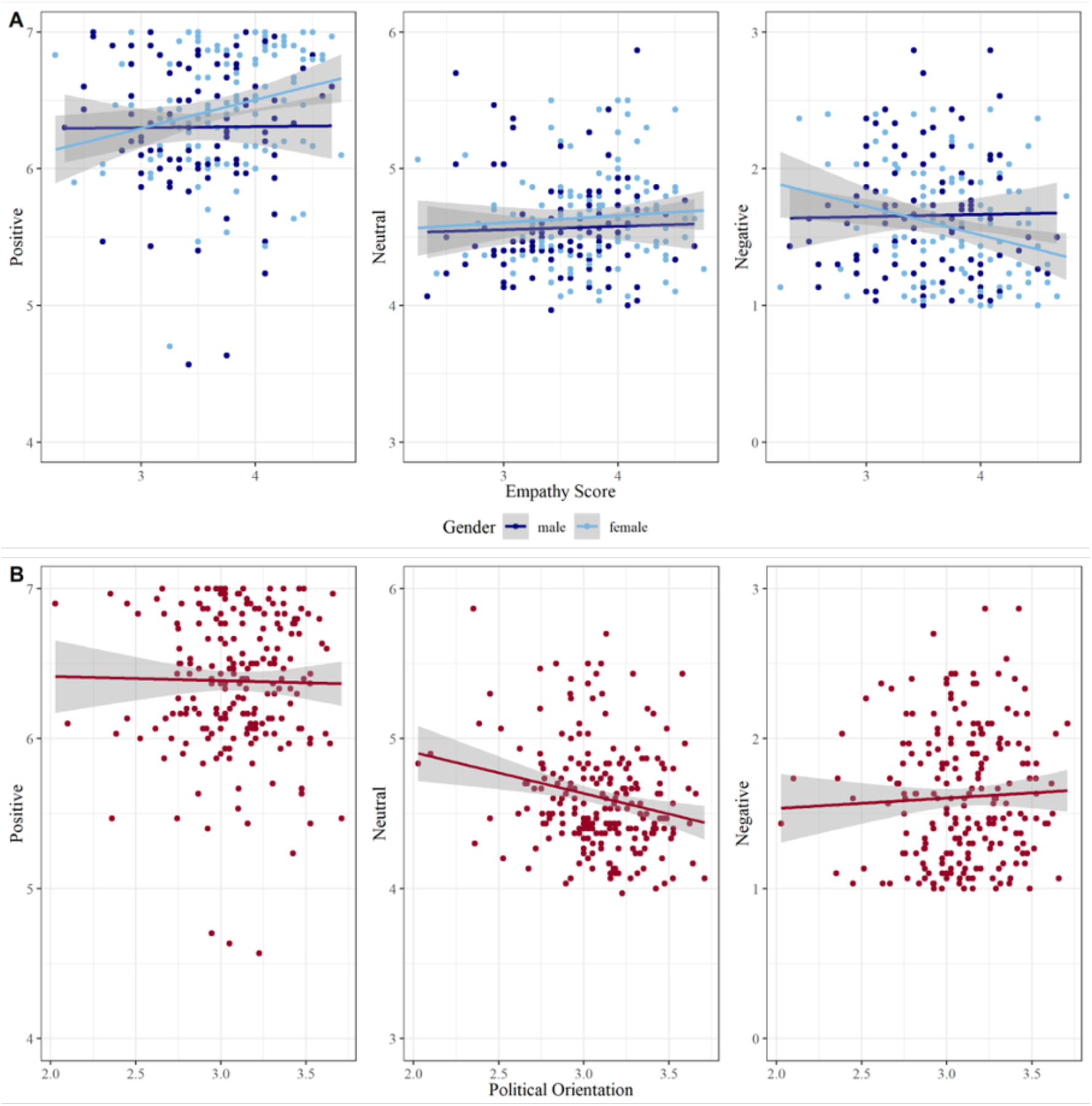
Influence of (A) the gender × empathy interaction and (B) political orientation on morality ratings. (A) Scatterplots show morality ratings as a function of empathy (1 = low empathy, 5 = high empathy) and gender across the three moral valence conditions (positive, neutral, negative). (B) Scatterplots depict morality ratings as a function of political orientation (1 = very conservative, 4 = very progressive) across the three valence conditions.

### Valence-specific models

- **Helpful**: Overall model *χ²*(4) = 12.68, *p* = .013, *R²_marginal_* = 0.02. Gender predicted ratings (*t* = 2.05, *p* = .040), with females rating helpful acts more positively; no effects of empathy, political orientation, or their interaction.
- **Harmful**: Overall model *χ²*(4) = 10.88, *p* = .028, *R²_marginal_* = 0.01. Gender × empathy interaction (*t* = -1.99, *p* = .047): higher empathy predicted harsher judgments among women only.
- **Neutral**: Overall model *χ²*(4) = 14.66, *p* = .005, *R²_marginal_* = 0.01. Political orientation (*t* = -3.66, *p* < .001): progressives rated neutral acts less positively than conservatives; no other significant effects.

See Appendix, Table 2 for complete results.

**Table 2.** Example of one moral scenario in three versions for each moral valence (negative, neutral, positive)

| moral category | scenario |
| --- | --- |
| negative | Peter and Anna take the train to Frankfurt and collide at the door of a compartment. Both have the same seat number on their reservation due to a booking error. Peter insists on his seat, although Anna has a broken leg. |
| neutral | Peter and Anna take the train to Frankfurt and collide at the door of a compartment. Both have the same seat number on their reservation due to a booking error. Peter leaves the seat to Anna and takes another free seat. |
| positive | Peter and Anna take the train to Frankfurt and collide at the door of a compartment. Both have the same seat number on their reservation due to a booking error. Peter leaves the seat to Anna and sits down in the aisle. |

*Note.* 180 scenarios (each 60 negative, neutral, positive) were presented for each participant. Each participant was shown only one out of the three scenarios for each setting.

### Correlation Between Empathy and Political Orientation

Empathy correlated positively with progressive political orientation, *r* = .22, *p* = .001, 95% CI [0.09, 0.34].

## Interim Discussion

Study 1 investigated how empathy, political orientation, and gender influence moral judgments of unfamiliar faces, embedded in brief, face-paired scenarios. Moral impressions were influenced both by the valence of the described act and, to lesser extent, by stable individual differences.

As expected, participants rated helpful agents most positively, harmful agents most negatively, and neutral agents in between, supporting the ecological validity of the paradigm (Uhlmann et al., 2015). Neutral acts received mildly positive ratings, suggesting a “benign default” toward norm-conforming behavior (Sears, 1983), alongside a negativity bias for harmful acts.

Gender and empathy interacted: more empathic women judged harmful acts more harshly, consistent with greater care orientation in women (Jaffee & Hyde, 2000), This suggests that empathy may heighten emotional sensitivity to norm violations, especially in female observers. Given the small effect sizes, these findings should be interpreted cautiously, as meta-analyses indicate that gender differences in moral reasoning are typically modest and context dependent (Hyde, 2005).

Contrary to our hypothesis, political orientation affected only judgments of neutral scenarios: conservatives rated neutral behavior more positively than progressives. While conservatism has been linked to greater reactivity toward moral violations (Hibbing et al., 2014; Oxley et al., 2008), our findings suggest conservatives may also assign greater moral value to norm-conforming or ambiguous behavior. This interpretation is consistent with evidence that conservatives more often interpret socially expected behavior as norm-reinforcing or virtuous (Jost et al., 2003; Joel et al., 2014). Empathy and political orientation were positively correlated, consistent with reports that progressives tend to report higher empathic concern (Hasson et al., 2018; Waytz et al., 2016).

Our paradigm’s use of ecologically valid moral scenarios – including helpful and neutral behavior as well as transgressions – offers a broader, more naturalistic basis for person-centered moral judgment than stylized dilemmas (Kahane, 2015), By presenting a broader range of everyday actions, our paradigm bridges the gap between abstract moral reasoning and real-world impression formation (Knutson et al., 2010). This aligns with calls to capture the full moral spectrum for a comprehensive understanding of moral cognition (Hart et al., 2024).

Limitations of our study include the small size of empathy and gender effects, a predominantly student sample limiting demographic diversity, and reliance on self-report measures that may not capture the multidimensionality of empathy and political orientation. Future research should examine other moral domains (e.g., purity, fairness; Graham et al., 2013) and contexts varying in relational closeness. Study 2 addresses some of these issues by recruiting ideologically more distinct participants, by applying EEG to track temporal dynamics, and by extending scenarios to varied moral foundations.

In sum, even brief contextual information about social behavior can shape moral impressions of unfamiliar individuals. These impressions vary with behavior valence and observer traits – particularly empathy, gender, and, for ambiguous acts, political orientation – underscoring the value of a person-centered approach by combining ecologically valid materials with rigorous quantitative modeling.

### Study 2 – Neural responses to morally contextualized faces

Study 2 investigated whether the person-centered moral evaluations observed in Study 1 are also reflected in neural processing, and whether these effects vary by political orientation. Using an EEG adaptation of the face-scenario paradigm, we paired neutral faces with descriptions of helpful, harmful or neutral behavior. Such associative learning approaches have been shown to reliably modulate neural responses to inherently neutral faces (e.g., Hammerschmidt et al., 2017; Hammerschmidt et al., 2018; Suess et al., 2014; Wieser et al., 2014; Ziereis & Schacht, 2023). Previous research suggests that even a single piece of moral information can alter attentional and evaluative processes of faces, as reflected in ERP components such as the P1, N170, Early Posterior Negativity (EPN), and Late Positive Potential (LPP) (e.g., Baum et al., 2018; Huang et al., 2024; Leuthold et al., 2014; Zhang et al., 2025).

Unlike Study 1, which focused on explicit judgments of moral behavior, Study 2 used likeability ratings as a more affective and socially grounded measure suited to ERP methodology, which captures rapid, automatic processes preceding deliberate reasoning. We focused solely on the agent’s face, as Study 1 showed that agent behavior alone shaped impressions. This allowed us to isolate neural correlates of moral person knowledge without the complexity of a dyadic agent– patient structure.

To examine individual differences, we recruited participants from a large online prescreening sample of over 600 individuals, who scored at the conservative or progressive extremes on a validated political orientation scale (Demel et al., 2023).

## Hypotheses

Based on the literature described above, we preregistered the following hypotheses (https://osf.io/q6fzy/?view_only=166764ff59af494da8e860029d88acc9):

1. **Likeability ratings:** Moral context will influence ratings, with negative actions leading to lower likeability ratings than neutral actions and positive actions leading to higher likeability ratings than neutral actions.
2. **ERP amplitudes (LPP):** Faces associated with negative actions will elicit enhanced LPPs compared to those linked with neutral or positive actions, reflecting greater affective salience.
3. **Political orientation – behavior:** Conservatives will rate negative agents less likeable and positive agents more likeable than progressives.
4. **Political orientation – ERP:** Political orientation will moderate LPP effects, with progressives showing stronger differentiation between moral contexts than conservatives. Earlier components (P100, N170, EPN) may also be modulated, particularly for faces linked to harmful behavior, consistent with rapid perceptual and attentional sensitivity to negative moral cues.

## Methods

### Participants

We conducted a large-scale pre-screening with 675 participants (455 female, 216 male, 4 other; *M_age_* = 24.63, *SD* = 5.76) using an early version of the Contemporary German Questionnaire of Political Orientation (CGPOQ; Demel et al., 2023), along with demographic and empathy questionnaires (see Study 1).

Those scoring at least ±0.75 SD from the group mean of the CGPOQ validation sample were invited to the EEG study, yielding two matched groups with divergent political orientations: 20 progressive and 20 conservative participants (20 female, 20 male; *M_age_* = 25.32, *SD* = 6.25, range = 18–44). All participants were German native speakers, right-handed (Oldfield, 1971), and reported no neurological or psychiatric disorders. One conservative participant was excluded due to EEG data loss >50%, leaving *N* = 39 for analysis.

### Sample Size Rationale

Sample size was determined via power simulation in R (R Core Team, 2018), targeting a small effect size (*f* = .10), a cumulative Type I error rate of 5%, and within-subject intercorrelations of *r* = .70 based on prior P100 effects (Demel et al., 2019). A pre-registered group-sequential design with interim stopping rules (https://osf.io/q6fzy/?view_only=166764ff59af494da8e860029d88acc9) set a maximum of *N* = 80. Recruitment ended at 40 completed datasets due to COVID-19 and difficulty recruiting conservatives, still exceeding sample sizes of similar ERP studies (e.g., Hammerschmidt et al., 2017).

### Ethics

The study was approved by the ethics committee of the Institute of Psychology, University of Göttingen, and conducted in accordance with the Declaration of Helsinki. Participants received either course credit or monetary compensation.

### Design and Procedure

Participants completed an EEG adaptation of the face–scenario task from Study 1. After providing written informed consent, they completed demographic questions, a left–right political orientation self-rating (1 = left, 7 = right), and the Edinburgh Handedness Inventory (Oldfield, 1971). Participants were seated in a sound-attenuated, dimly lit booth, facing a monitor (3544 × 6260 px) at a distance of 77 cm. EEG electrodes were applied, heads were stabilized in a chinrest. After instructions they completed one practice trial with stimuli not included in the main task.

Each trial began with a black fixation cross (54 px; 1,000 ms), followed by a neutral face (270 × 390 px, visual angle ∼5.3° × 7.7°; 1,000 ms), and a randomly assigned name label (3,000 ms). A moral scenario was then presented for at least 7,000 ms, with the agent’s face above the text. The face-name-scenario combinations were gender-matched and randomized across trials. Participants could advance via mouse click, or the screen advanced automatically after 15,000 ms. After a jittered fixation (500–1,000 ms), the same agent face reappeared for 1,000 ms. ERPs were time-locked to this face presentation. Following a fixation (1,000 ms), participants rated the likability (–3 = very unlikable to +3 = very likable), with the face still displayed above the scale (see Figure 3). The experiment was programmed using *PyCharm* (JetBrains, 2017) and implemented in Python.

**Figure 3.**
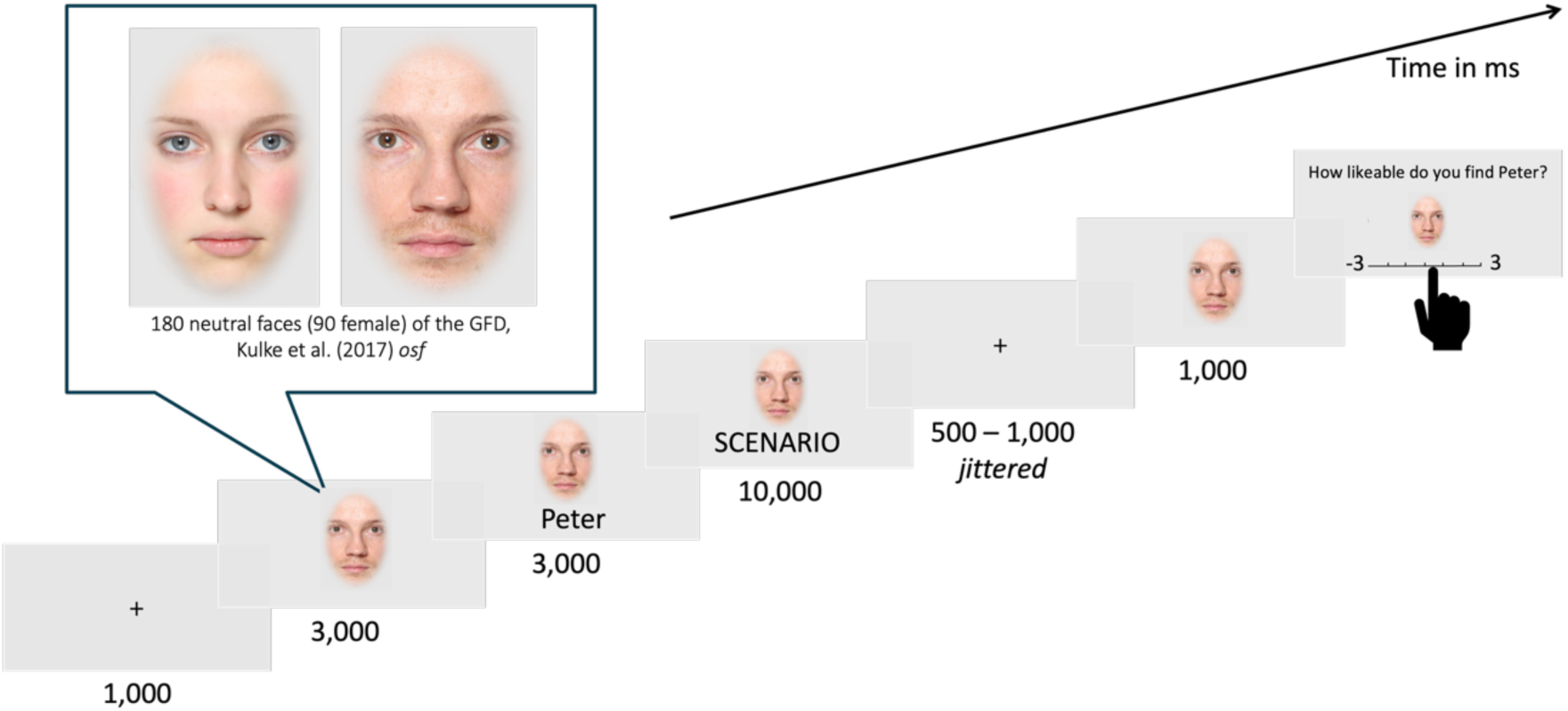
Trial Scheme of Study 2. In total, 180 trials were presented with a full randomization of face, scenarios, name, and presentation order.

Participants completed 180 unique trials (10 blocks of 18 sccenarios), with all faces, names, scenarios presented once. Short breaks (1-5 minutes) were allowed between blocks. Including EEG setup, sessions lasted ∼2.5-3 hours.

### Stimuli

180 mixed gender scenarios (60 per moral valence) were drawn from a validated stimulus set of 540 scenarios where a moral agent commits either a harmful, helpful, or neutral act towards a patient. The scenarios were based on all five moral foundations (Graham et al., 2013). Furthermore, they exist in three parallel versions which means that the setting of the scenario is always the same but differs in the action from the agent towards the patient. Each participant saw only one of the parallel versions (see Table 2, for an example). Faces were selected from the Göttingen Face Database (Kulke et al., 2017) and pre-rated for neutral expression. Each face–name–scenario combination was unique and gender-balanced.

### EEG Recording and Preprocessing

EEG was recorded from 64 Ag/AgCl electrodes (Easy-Cap; Biosemi, Amsterdam, The Netherlands) with Common Mode Sense (CMS) and Driven Right Leg (DRL) as reference and ground (cf. http://www.biosemi.com/faq/cms&drl.htm). Two external electrodes were placed on the left and right mastoids (A1, A2), and four EOG electrodes (vertical and horizontal) monitored eye movements.

EEG was sampled at 512 Hz, bandpass-filtered online (0.16–100 Hz), and electrode offsets kept within ±20 μV.

Preprocessing in Brain Vision Analyzer (Brain Products GmbH, Munich, Germany) included downsampling to 500 Hz, re-referencing to average reference, and offline filtering (low cutoff of 0.03 Hz, time constant: 5 s; 12 dB/oct; high cutoff of 40 Hz, 48 dB/oct) and a 50 Hz notch filter.

Epochs were extracted from –200 to 900 ms relative to face stimulus onset, with a 200 ms pre-stimulus baseline. Blink artifacts were corrected using the Gratton– Coles method (Gratton et al., 1983). Artifact-contaminated segments were excluded, and remaining epochs were averaged for each participant and condition (harmful, neutral, helpful agent).

ERP components and their corresponding regions of interest (ROIs) were defined per preregistration and refined via visual inspection of aggregate grand averages (AGATs) as recommended by Brooks et al. (2017):

- **P100**: Most positive peak between 80–120 ms, referenced to O1. ROI included O1, O2, PO7, PO8, Iz, Oz to better capture the widespread occipital positivity observed across conditions.
- **N170**: Most negative peak between 140–176 ms, referenced to P9. ROI included standard parieto-temporal electrodes: P7, P8, P9, P10, PO7, PO8, TP7, TP8.
- **EPN (Early Posterior Negativity)**: Mean amplitude between 200–350 ms. ROI based on preregistration but adjusted following group-level inspection of difference waves (progressive vs. conservative): P5, P6, PO7, PO8, P7, P8.
- **LPP (Late Positive Potential)**: Mean amplitude between 350–600 ms. ROI was adapted to reflect a more occipital distribution of the signal than initially preregistered: O1, O2, PO3, PO4, PO7, PO8, Oz.

All preprocessing choices and adjustments were made prior to statistical analysis and transparently according to open science practices. Data and analysis script can be found under https://osf.io/acmkr/?view_only=cdf220d0dbc24114b46f1b012c6b8a7f.

## Results

### Sample Characteristics and Political Orientation

In the prescreening (N = 675), the sample leaned progressive (*M* = 36.43 on a 0–100 scale; 0 = very progressive, 100 = very conservative). Empathy and political orientation were significantly correlated (*r* = –.55, *R²* = .30, *p* < .001, 95% *CI* [0.23, 0.37]), with progressives reporting higher empathy.

For EEG testing, 20 high-conservative and 20 high-progressive participants were selected (at least ±0.75 SD in CGPOQ), matched for age and gender: (10 females and 10 males). Conservatives were slightly older (*M_age_* = 26.05, *SD* = 6.89) than progressives (*M_age_* = 23.75, *SD* = 3.89). Group differences were robust for validated CGPOQ scores (*t*(36.08) = 26.18, *p* < .001) and left-right self-placement (*t*(36.1) = 9.93, *p* < .001), confirming successful stratification. No group differences emerged in self-reported empathy (*t*(37.24) = –1.19, *p* = .244) or religiousness (*t*(37.13) = 1.38, *p* = .177).

### Behavioral Data: Likeability Judgments

Likeability ratings (–3 = very unlikeable, +3 = very likeable) varied by scenario valence: harmful agents were rated negatively (*M* = –2.04, *SD* = 1.07), neutral agents moderately (*M* = 0.79, *SD* = 1.22), and helpful agents positively (*M* = 1.69, *SD* = 1.12). This pattern was consistent across conservatives and progressives.

A linear mixed-effects model (fixed effects: valence, political orientation, agent gender, and their interactions; random intercepts for participants and items) showed a strong main effect of valence on likeability judgments, *χ*²(6) = 8099.49, *p* < .001, *R²_marginal_* = 0.66. Planned comparisons confirmed more positive ratings for helpful vs. neutral agents (*p* < .001) and more negative ratings for harmful vs. neutral agents (*p* < .001). No interactions with political orientation or agent gender were significant.

### ERP Results

**P100 and N170:** There were no significant effects of moral valence or political orientation, indicating that prior moral information did not influence early perceptual encoding.

**EPN (200–350 ms):** There was no modulation by valence or political orientation, suggesting moral context did not affect emotion-related attention.

**LPP (350–600 ms):** Amplitudes were significantly modulated by moral valence, *χ2* = 11.43, *df* = 4, *p* = .022, and by an interaction between moral valence and political orientation, *χ2* = 6.31, *df* = 2, *p* = .047, *R^2^_marginal_* = 0.01. Positive agents elicited higher amplitudes than negative agents (*p* = .021), driven by progressives (positive > negative: *p* = .003) but not conservatives (*p* = .573). The neutral–positive difference was larger for progressives than conservatives (*p* = .020); for conservatives there was no significant difference, whereas progressives showed larger amplitudes for positive vs. neutral agents (see Figure 4).

**Figure 4.**
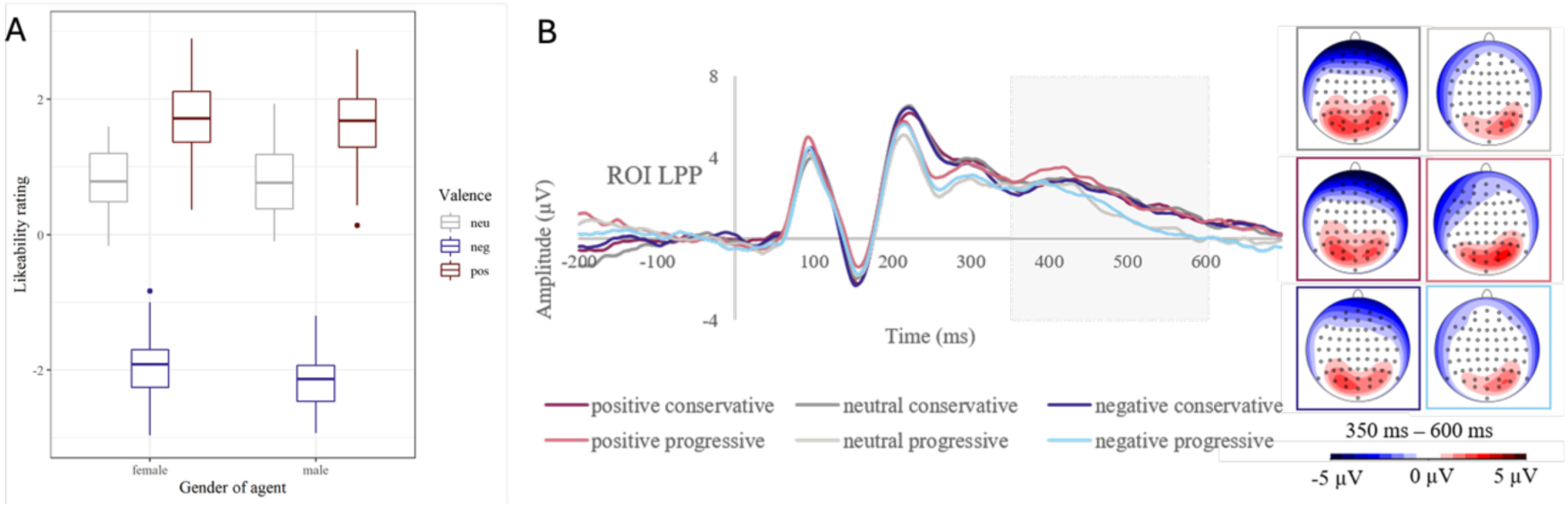
Main Behavioral and Neural Findings of Study 2. (A) Boxplots of likeability ratings as a function of agent gender and moral valence of the associated behavior. Agents described as helpful were rated most likeable, harmful agents least likeable, with male harmful agents rated slightly less favorably than female ones. (B) Grand average ERP waveforms (center) and topographies (right) of the LPP (350–600 ms) as a function of scenario valence (positive, neutral, negative) and political orientation (progressive vs. conservative). While conservatives showed generally elevated LPP amplitudes across conditions, progressives displayed enhanced LPPs selectively for faces associated with morally positive actions. Topographies display the difference waves (positive minus neutral [top], positive minus negative [middle], negative minus neutral [bottom]) separately for each group (left: conservatives; right: progressives). Color bar indicates voltage in microvolts (μV).

## Interim Discussion

Study 2 examined whether prior moral information influences likeability judgments and neural responses to faces, and whether these effects are moderated by political orientation. Consistent with hypothesis 1, agents described as acting immorally were rated as less likeable than those whose actions were neutral or helpful. Hypothesis 2 was partly supported: moral valence significantly modulated LPP amplitudes, but contrary to our expectation of a negativity bias, positive (vs. negative) agents elicited higher amplitudes. This aligns with evidence that morality is central to impression updating and person perception (Brambilla et al., 2019; Brambilla & Leach, 2014). For example, Singer et al. (2004) showed that faces linked to fair versus unfair behavior modulated both affective evaluations and neural responses related to empathy and reward processing. Interestingly, we did not observe the typical negativity bias in the LPP reported in prior studies (Bartholow et al., 2001; Hajcak & Olvet, 2008; Weinberg & Hajcak, 2010). Instead, helpful agents elicited the strongest LPP responses. This pattern may be due to the structure of our scenarios, many of which began with a negative situation that was resolved through prosocial behavior. Such an expectation-violating shift from negative to positive likely requires greater cognitive elaboration, thereby enhancing the LPP (Du et al., 2014; Bartholow et al., 2001). Positive moral behavior, though ecologically relevant, remains underrepresented in ERP research, and our results underscore the need for further investigations of such behavior (Yoder & Decety, 2014).

Contrary to Hypothesis 3, political orientation did not influence likeability ratings, with conservatives and progressives evaluating agents similarly. This contrasts with prior work linking conservatism to heightened moral sensitivity and emotional reactivity (Hibbing et al., 2014; Oxley et al., 2008; Jost et al., 2003). Likeability judgments may capture affective resonance rather than moral evaluation per se, and strong ceiling/floor effects in our clearly valenced scenarios may have masked subtler ideological differences. More nuanced measures (e.g., moral responsibility, trustworthiness, or willingness to interact) could offer greater sensitivity. Large-scale data provide further context: Casey et al. (2023) found that liberals express less empathy toward political out-group members, partly via harsher moral judgments, and Hart et al. (2024) proposed the asymmetry hypothesis: binding moral values, often endorsed by conservatives, more strongly predict endorsement of virtuous behavior than condemnation of transgressions. These findings suggest that ideological differences may emerge more clearly in evaluations of ambiguous or norm-conforming acts rather than in overt moral violations.

Hypothesis 4 was supported: conservatives showed consistently elevated LPP amplitudes across all moral conditions, suggesting a generally heightened affective response to morally contextualized faces. This pattern aligns with previous findings indicating that conservatives exhibit enhanced LPPs and heightened reactivity to low-arousing stimuli (Tritt et al., 2016); it extends these findings by showing that such physiological differences also occur for moral information. Progressives, by contrast, displayed selective LPP responses for helpful agents, indicating greater sensitivity to positive moral valence.

The divergence between similar likeability ratings and differing neural responses may point to distinct processing styles: conservatives may rely more on rapid, intuitive (System I) processing, whereas progressives engage on slower, deliberative (System II) evaluation (Lane & Sulikowski, 2017). Faster, more emotional responses among conservatives (Tritt et al., 2013) may underlie the uniformly high LPP amplitudes observed.

The final sample was smaller than planned in the pre-registration, which may have limited power to detect subtle effects. However, the study is still comparable to or larger than many similar ERP studies (e.g., Hammerschmidt et al., 2017; Leuthold et al., 2014; Tritt et al., 2016).

## General Discussion and Conclusion

The present research examined how individual differences – particularly empathy, political orientation, and gender – influence moral impression formation, and whether these effects extend from explicit evaluations to neural processing of faces. Across two studies, we combined a naturalistic moral scenario paradigm with behavioral ratings (Study 1, Study 2) and EEG measures (Study 2) to capture both conscious judgments and affective–motivational processes. By integrating person-related traits with moral context manipulations, our aim was to clarify how stable dispositions shape responses to morally relevant behaviors, and to assess whether these influences emerge consistently across behavioral and neural levels.

## Integration and Implications

Together, Studies 1 and 2 provide a complementary view of how person-related and contextual variables shape moral evaluation at behavioral and neural levels. Both used a carefully constructed and validated set of moral scenarios embedded in a person-centered paradigm, pairing neutral faces and names with helpful, harmful, or neutral acts. This approach addresses calls for greater ecological and social validity in moral psychology (Hester & Gray, 2020; Uhlmann et al., 2015) by situating judgments within identifiable social targets.

Across studies, moral valence robustly influenced evaluations, with helpful agents viewed most positively and harmful agents most negatively. Person-level characteristics shaped these impressions in distinct ways: in Study 1, empathy and gender interacted in responses to harmful behavior, and political orientation affected ratings only for neutral acts – suggesting ideological differences emerge more in morally ambiguous contexts. In both samples, empathy correlated positively with progressive orientation, consistent with links between socio-affective dispositions and political values (Hasson et al., 2018; Waytz et al., 2016). Study 2 extended these findings to neural processing: conservatives showed consistently elevated LPP amplitudes across conditions, while progressives exhibited selective enhancement for helpful agents. This dissociation between explicit and neural measures suggests that ideological differences in motivational salience can occur without overt rating differences, aligning with evidence for heightened affective reactivity in conservatives (Oxley et al., 2008; Tritt et al., 2016) and dual-process accounts of moral judgment (Greene et al., 2001; Lane & Sulikowski, 2017). The divergence between similar ratings and differing LPP patterns underscores the value of combining explicit and neural measures to capture ideological differences that may not be evident behaviorally.

## Limitations and Future Directions

Our studies provide novel insights into moral evaluation across behavioral and neural levels, but several limitations remain. First, the two studies differed in task framing, measures, and sample composition, limiting direct comparability; aligning key design parameters would enable more direct cross-modal inferences.

Second, although our EEG sample was relatively large for ERP research, it was smaller than intended potentially reducing sensitivity to subtle effects. Future replication with larger, more diverse samples could confirm the observed interaction patterns and extend them to other traits (e.g., religiosity, moral foundations).

Third, our stimuli – neutral faces with brief scenario vignettes – were systematically constructed and validated, allowing for tight control but may not capture the richness of real-world moral evaluation. More ecologically valid contexts, such as dynamic video scenarios, social media profiles, or immersive virtual reality, could reveal additional facets of moral cognition.

Fourth, political orientation in Study 2 was dichotomized into progressives versus conservatives to maximize contrast, but this oversimplifies multidimensional construct. Continuous, multidimensional measures (e.g., Demel et al., 2023) and analytic approaches like cluster analysis could capture more nuanced ideological effects.

Finally, empathy – linked to political orientation in both samples – was not included as a predictor in the EEG study because of the small sample size, limiting conclusions about its role in neural responses. Incorporating empathy across behavioral and neural paradigms in large samples could clarify whether it mediates or moderates ideological differences. The divergence between behavioral and ERP findings further underscores the need to examine how conscious and implicit processes interact in moral judgment.

By increasing sample sizes, aligning experimental designs, and enhancing ecological validity, future research can build on these findings to advance a socially grounded model of moral cognition.

## Conclusion

Together, our two studies demonstrate that moral evaluation is shaped by the interplay of stimulus features, perceiver traits, and processing level. By combining large-scale behavioral data with high-temporal-resolution EEG measures, we show that ideological differences in motivational salience can emerge without overt rating differences. These findings highlight the importance of incorporating both morally positive and negative behaviors in ecologically valid, person-centered paradigms, and of integrating behavioral and neural approaches to capture the full complexity of moral cognition.

## Acknowledgments

The authors thank Thomas Schultze-Gerlach for statistical support and Jana Thiel for help with data collection.

## CRediT Authorship Contribution Statement

RD: Conceptualization, Funding Acquisition, Methodology, Data curation, Formal analysis, Visualization, Writing; MRW: Conceptualization, Funding Acquisition, Writing, Supervision; AS: Conceptualization, Funding Acquisition, Resources, Methodology, Validation, Writing, Supervision.

## Conflicts of interest

All authors declare no conflicts of interest.

## Funding

This research was supported by the Stiftung der Deutschen Wirtschaft, a Leibniz ScienceCampus Primate Cognition grant, and the RTG 2070 “Understanding Social Relationships”, the CRC 1528 “Cognition of Interaction”, and a Reinhart Koselleck project grant (WA 621/25-1), funded by the Deutsche Forschungsgemeinschaft (DFG).

***Transparency and Openness.*** All data files, and analysis scripts are available under https://osf.io/qsbvx/?view_only=dd12d36861f240288d3533724fcce665 (Study 1), preregistration is available under https://osf.io/q6fzy/?view_only=166764ff59af494da8e860029d88acc9 and data files and analysis script are available under https://osf.io/acmkr/?view_only=cdf220d0dbc24114b46f1b012c6b8a7f (Study 2). All further materials as well as raw EEG data can be made available upon request.

## Appendix

**Table 1.**
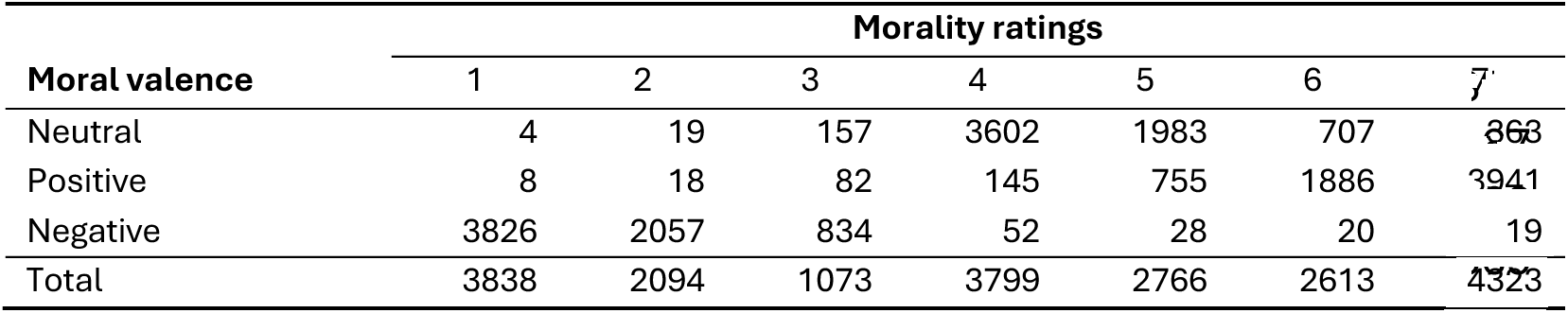
Distribution of morality ratings for each moral valence.

**Table 2.** Main effects and interactions for the three models of each moral valence.

|  | Likelihood Ratio Tests |  |  |  |  |  |
| --- | --- | --- | --- | --- | --- | --- |
|  | Estimate | SE | t | p | CI<sub>lower</sub> | CI<sub>upper</sub> |
| **Helpful(pos)** |  |  |  |  |  |  |
| intercept | 6.30 | 0.07 | 93.71 |  | 6.169 | 6.436 |
| empathy<sup>(1)</sup> | 0.01 | 0.05 | 0.19 | .849 | -0.824 | 0.100 |
| gender | 0.13 | 0.06 | 2.05 | **.040** | 0.005 | 0.253 |
| political orientation<sup>(1)</sup> | -0.03 | 0.03 | -0.98 | .332 | -0.092 | 0.031 |
| gender x empathy | 0.10 | 0.06 | 1.67 | .096 | -0.019 | 0.229 |
| **Harmful(neg)** |  |  |  |  |  |  |
| intercept | 1.66 | 0.07 | 25.18 |  | 1.531 | 1.793 |
| empathy<sup>(1)</sup> | 0.00 | 0.04 | 0.04 | .968 | -0.085 | 0.088 |
| gender | -0.07 | 0.06 | -1.22 | .222 | -0.191 | 0.045 |
| political orientation<sup>(1)</sup> | 0.04 | 0.03 | 1.32 | .190 | -0.019 | 0.098 |
| gender x empathy | -0.12 | 0.06 | -1.99 | **.047** | -0.237 | -0.011 |
| **Neutral** |  |  |  |  |  |  |
| intercept | 4.58 | 0.10 | 45.87 |  | 4.384 | 4.786 |
| empathy<sup>(1)</sup> | 0.03 | 0.04 | 0.71 | .474 | -0.049 | 0.104 |
| gender | 0.06 | 0.05 | 1.10 | .271 | -0.046 | 0.161 |
| political orientation<sup>(1)</sup> | -0.96 | 0.03 | -3.66 | **< .001** | -0.147 | -0.044 |
| gender x empathy | 0.02 | 0.05 | 0.42 | .674 | -0.081 | 0.126 |

*Note.* <sup>(1)</sup> empathy and political orientation were z-transformed to a $M = 0$ and $SD = 1$ ; original means ( $SD$ ) were: empathy = 3.64 (0.51); political orientation: 3.10 (0.29).

## Notes

### Competing Interest Statement

The authors have declared no competing interest.

